# Competitive learning modulates memory consolidation during sleep

**DOI:** 10.1101/196964

**Authors:** James W. Antony, Larry Y. Cheng, Paula Pacheco, Ken A. Paller, Kenneth A. Norman

## Abstract

Competition between memories can cause weakening of those memories. Here we investigated memory competition during sleep by presenting auditory cues that had been linked to two distinct picture-location pairs during wake. We manipulated competition during learning by requiring subjects to rehearse item pairs associated with the same sound either competitively (choosing to rehearse one over the other, leading to greater competition) or separately; we hypothesized that greater competition during learning would lead to greater competition when memories were cued during sleep. With separate-pair learning, we found that cueing benefited spatial retention. With competitive-pair learning, no benefit of cueing was observed on retention, but cueing impaired retention of well-learned pairs (where we expected strong competition). During sleep, post-cue beta power (16-30 Hz) indexed competition-based weakening and forgetting, whereas sigma power (11-16 Hz) indexed memory strengthening. These findings show that memory consolidation during sleep fundamentally engages competition and selective memory weakening.

---

Despite the difficulty of precisely characterizing memory storage in the human brain, it is safe to assume that memories do not exist in a vacuum. Rather, each memory exists in a network with related memories. Moreover, it is rarely the case that a cue will evoke just one memory; instead, memories compete at retrieval, and this competition has consequences. Generally, when memories compete, winners are strengthened and losers are weakened (Anderson, Bjork, & Bjork, 2000; Norman, Newman, & Detre, 2007). For example, in retrieval-induced forgetting, recalling a subset of items within a category results in improvement for practiced items and impairment for non-practiced, related items within the same category (Anderson et al., 2000). On the other hand, several other studies have found that activating memories moderately (i.e., enough to compete, but not enough to win the competition) leads to their weakening (Detre, Natarajan, Gershman, & Norman, 2013; Kim, Lewis-Peacock, Norman, & Turk-Browne, 2014; Lewis-Peacock & Norman, 2014; Newman & Norman, 2010). These modifications adaptively shape the memory landscape, such that later retrieval is associated with reduced competition (Norman et al., 2007).

While the aforementioned studies of memory competition focused on wake learning, it is plausible that similar competitive dynamics could occur during sleep, with similarly important consequences. However, most studies of sleep and learning, with some notable exceptions (Genzel et al., 2017; Oyarzún, Moris, Luque, Diego-Balaguer, & Fuentemilla, 2017; Payne, Stickgold, Swanberg, & Kensinger, 2008), have focused on how sleep affects individual memories and not on how memories compete and the consequences of competition.

A substantial body of evidence shows that memories can be reactivated during sleep, leading to their stabilization (e.g., Gulati, Ramanathan, Wong, & Ganguly, 2014; Ji & Wilson, 2007; Wilson & McNaughton, 1994). Reactivation can be systematically biased when learning-related stimuli are presented during post-training sleep, a methodology termed targeted memory reactivation or TMR (Creery, Oudiette, Antony, & Paller, 2014; Diekelmann, Büchel, Born, & Rasch, 2011; Oudiette, Antony, Creery, & Paller, 2013; Rasch, Büchel, Gais, & Born, 2007; Rudoy, Voss, Westerberg, & Paller, 2009). Additionally, analyses of sleep electroencephalography (EEG) have linked memory strengthening via TMR with post-cue sigma power (Farthouat, Gilson, & Peigneux, 2017; Groch, Schreiner, Rasch, Huber, & Wilhelm, 2017; Lehmann, Schreiner, Seifritz, & Rasch, 2016; Schreiner, Lehmann, & Rasch, 2015). Similarly, memory weakening has been linked with post-cue beta power (Oyarzún et al., 2017).

In this study, we used TMR to study competition during sleep. We hypothesized that the degree of competition during sleep, along with the behavioral consequences of this competition, would depend on how memories are learned. Intuitively, the more that memories are entangled during wake, the more they will compete during sleep. We varied entanglement by either forcing memories linked by a common sound cue to be learned either in direct temporal proximity or separately. Moreover, competition can be enhanced if participants are forced to prioritize one memory over another, as opposed to allowing the memories to co-exist (Mather & Sutherland, 2011). We therefore explored the effects of prioritization by assigning a high or low reward to each item of a pair, and we included short periods allotted for memory rehearsal whereby subjects could prioritize encoding of high-reward items (Oudiette et al., 2013). Finally, we used TMR to bias reactivation of competing memories, and we hypothesized that increased competition would result in memory weakening.

Subjects (*N* = 60; Fig 1) first learned arbitrary associations between specific environmental sounds and visual items that were either celebrities, landmarks, or common objects. Some sounds were linked with two items from different categories (paired). For example, if a “meow” sound was presented with both Brad Pitt and the Eiffel Tower on two separate trials, then Brad Pitt and the Eiffel Tower became paired items via their mutual association to the sound “meow.” We included other sounds linked with one item (singular) to later assess the neural effects of simultaneously cueing one versus two memories during sleep. Next, subjects learned the spatial location of each item against a background grid during four rounds of encoding. At this point, each item was assigned either a high (9 cents) or low (1 cent) monetary reward that could be earned during subsequent memory tests. Singular items were shown alone and paired items were shown in groups of two (with a randomized order of presentation of one high-reward and one low-reward item) before a 5-s memory rehearsal period. Each item was presented with its accompanying sound. We further manipulated the level of competition between paired items (in a between-subjects fashion) by varying the temporal proximity of spatial location learning trials. Paired items were either presented in the same group in a competitive-pair learning condition (CPL; *n* = 30) or in different groups in a separate-pair learning condition (SPL; *n* = 30). For CPL, paired items were grouped together on all four rounds of encoding. For example, Brad Pitt’s location was always grouped with that of the Eiffel Tower. For SPL, paired items were never grouped together. Instead, items were grouped with different items of the opposite reward value on each round of encoding. For example, Brad Pitt might have been grouped with a globe on round one and the Taj Mahal on round two, etc, but never with the item that was also associated with the same “meow” sound (Eiffel Tower). In both cases, subjects were encouraged to prioritize location rehearsal for the high-reward item over that of the low-reward item.

**Figure 1.**
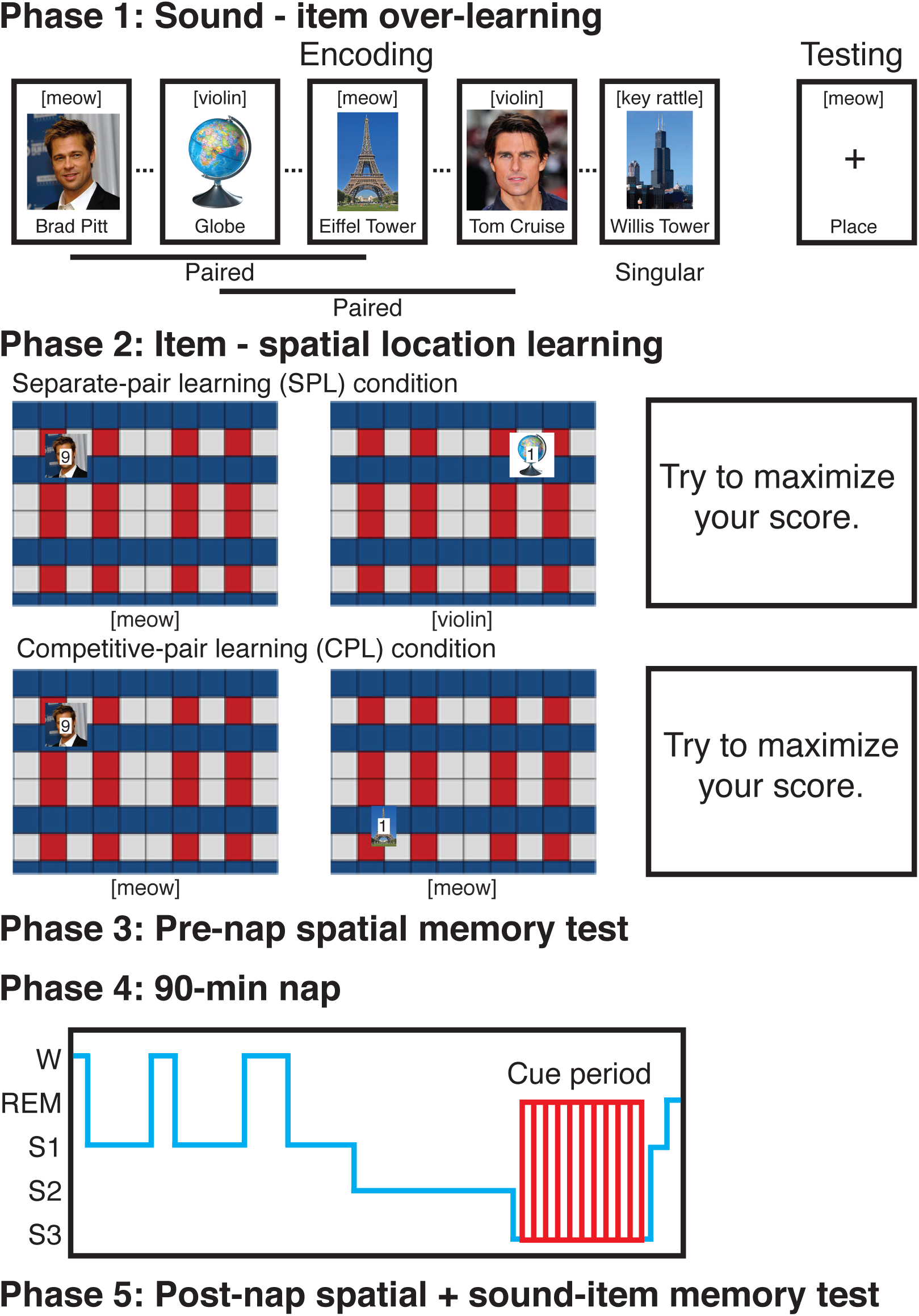
Task design. In phase 1, subjects over-learned associations between sounds and items. Associations were tested along with the category of the item until each association was correctly retrieved twice in a row. In phase 2, subjects encoded spatial locations for the same items (along with their accompanying sounds) against a background grid. Each paired item was assigned a high or low reward to be given upon correct recall, and each singular picture was assigned a high reward. The numbers depicting reward are enlarged here for expository purposes and did not obscure the item in the actual experiment. Paired items were shown in groups of two for 1-s each before a 5-s period when subjects were to prioritize rehearsal to maximize their score. Under separate-pair learning conditions (SPL), both items within a group were associated with different sounds, whereas under competitive-pair learning conditions (CPL), they were associated with the same sound. Singular pictures were shown alone before a similar 5-s prioritized recall period. In phase 3, subjects took a test on each spatial location. In phase 4, sounds from all singular and half of the paired items were presented during SWS. In phase 5, subjects took a final test on each spatial location followed by a sound-item test on the associations formed during phase 1.

Following this learning procedure, subjects were tested on their memory for item locations, and then took a nap in the lab. When intervals of slow-wave sleep were detected online by the experimenter, sound cues (100% of singular and 50% of paired sounds) were presented to the subjects while they slept. After the nap, subjects took a final item-location memory test and a sound-item association test. Subjects received an added monetary reward based on their performance on the pre-nap and post-nap item-location memory tests.

We predicted that the level of competition would impact TMR, such that SPL (low competition) and CPL (high competition) would lead to memory benefits and impairments, respectively. Furthermore, we predicted that paired low-reward items would be more susceptible to impairment than high-reward items. We also predicted (based on prior results reviewed above) that post-cue sigma power would positively predict subsequent memory and that post-cue beta power would index greater competition and negatively predict subsequent memory.

## Results

### Competition during learning impaired pre-nap accuracy

We first assessed whether competition affected learning prior to sleep. Our design included a between-subjects manipulation (CPL vs. SPL) and three item types: singular items (high-reward items associated with only a single sound), high-reward paired items, and low-reward paired items. We therefore submitted pre-nap spatial errors to a mixed, 2 (condition: CPL vs. SPL) × 3 (item type: singular, high-reward, or low-reward) ANOVA. We found a significant main effect of item type [*F*(2,116) = 59.2, *p* < 0.001], a marginally significant main effect of condition [*F*(1,58) = 3.2, *p* = 0.08], and a significant interaction [*F*(2,116) = 7.8, *p* < 0.001). As shown in Fig. 2A, follow-up *t*-tests revealed pre-nap recall accuracy was better in the SPL than for the CPL condition for singular items [SPL: 128.4 ± 12.7 pixels, CPL: 173 ± 13.4, *t*(58) = 2.4, *d* = 0.63, *p* = 0.02] and for high-reward items [SPL: 138.7 ± 12.2, CPL: 183.4 ± 14.0, *t*(58) = 2.4, *d* = 0.62, *p* = 0.02] but not for low-reward items [SPL: 214.4 ± 12.6, low reward CPL: 215.9 ± 13.9, *t*(58) = 0.08, *d* = 0.01, *p* = 0.94]. Whereas this effect shows competition had a meaningful impact on learning, we did not anticipate an effect on singular items. We speculate that pair encoding might have been more difficult in the CPL condition, so subjects may have occasionally rehearsed previously-shown paired items during rehearsal periods that followed singular items, thus weakening memory for singular items.

### Competition during learning influenced the effects of targeted memory reactivation

Our primary procedural manipulations were 1) altering the amount of competition between paired items during learning by either presenting them competitively (CPL condition) or separately (SPL condition), 2) administering TMR cues for only half of the pairs, thus creating cued and uncued conditions, and 3) manipulating reward for each paired item to be either high or low. Our primary dependent measure of forgetting across the nap was computed as post-nap error minus pre-nap error, after regressing out the effects of pre-nap error (see Methods; Fig S1). Greater positive values indicate more forgetting and therefore worse memory retention.

We first asked whether competition affected TMR efficacy by contrasting cued – uncued scores (the cueing effect) across conditions. We found the cueing effect was larger in the SPL condition than the CPL condition [in pixel mean ± SEM, SPL cued – uncued error: −14.9 ± 6.0, CPL cued – uncued error: 7.9 ± 7.3, *t*(59) = 3.0, *d* = 0.78, *p* = 0.004], demonstrating that competition decreases the efficacy of TMR (Fig 2B). Next, to ensure differences in TMR efficacy were not merely driven by pre-nap memory differences between the conditions, we randomly resampled subjects without replacement (*N*=18 - 28) from both the CPL and SPL groups to find instances in which there was no pre-nap difference in high reward (*t* < 0.5). We found the interaction (greater cueing effect in SPL than CPL) still held in each of 100 instances meeting this condition as determined by a negative *t* value in each (*t* mean = −2.26, standard deviation = 0.46, range = −3.65 to −1.2). Therefore, pre-nap memory differences between the conditions cannot explain the differences in TMR efficacy.

**Figure 2.**
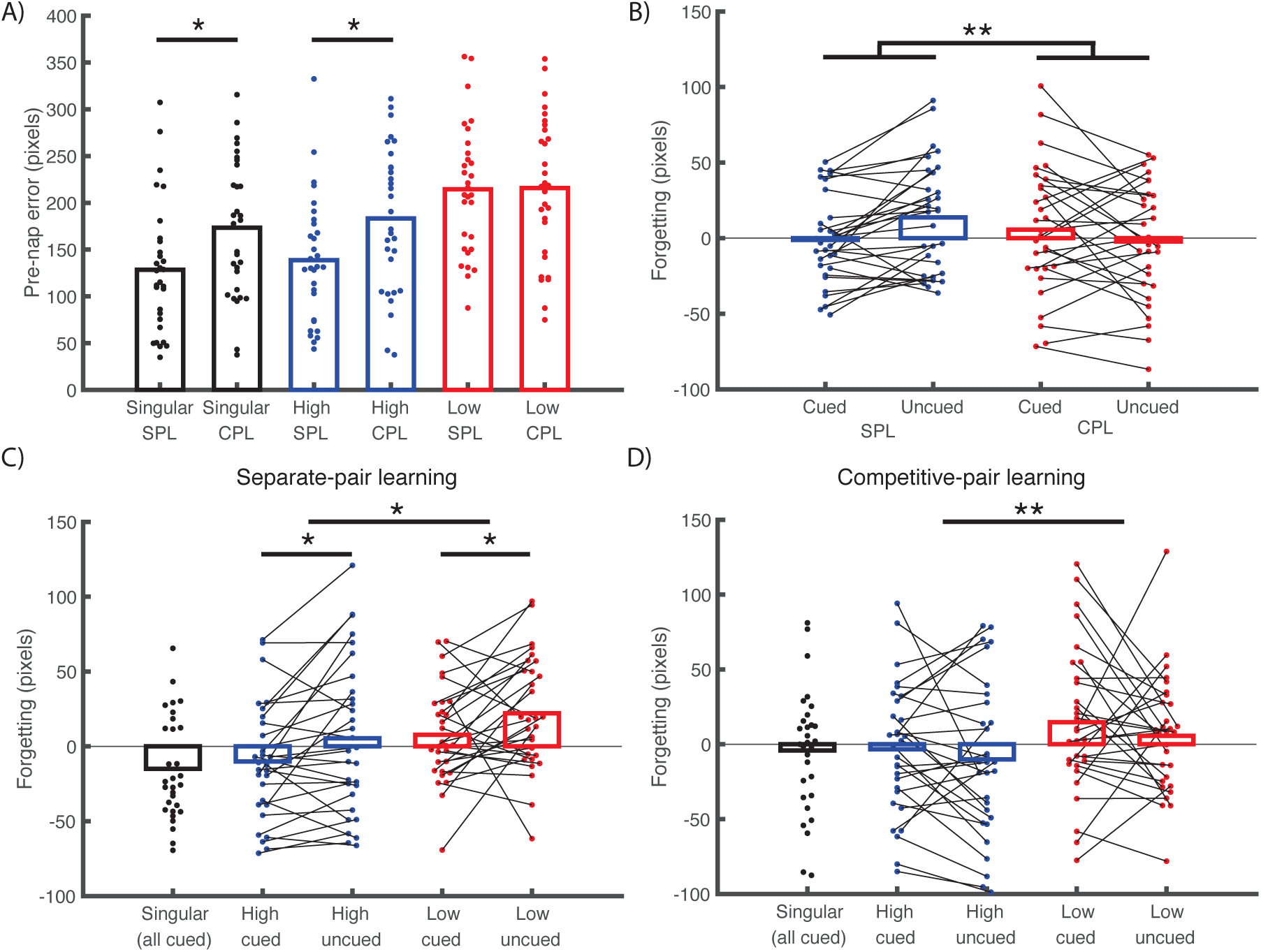
Inter-item competitive learning influences learning and targeted memory reactivation. We depicted data for various conditions using bee swarm plots, with rectangular box heights indicating means. (A) Competition negatively affected learning, as shown by pre-nap error between the conditions. (B) Competition strongly altered the effectiveness of targeted memory cues, where lower forgetting indicates better memory retention. (C) Under SPL, reward reduced forgetting, and cueing reduced forgetting for both high and low rewards. (D) Under CPL, reward reduced forgetting, but cueing had no overall significant effect on memory. *: *p* <= 0.05. **: *p* < 0.01.

Given that competition had a significant effect on TMR efficacy, we next assessed the effects of reward priorities and TMR on memory retention using a two-way, repeated measures ANOVA. For SPL, high-reward items were better remembered than low-reward items [*F*(1,29) = 6.0, *d*_*z*_ = 0.45, *p* = 0.02] and cued items were better remembered than uncued items [*F*(1,29) = 11.0, *d*_*z*_ = 0.61, *p* = 0.002], but there was no interaction between the conditions [*F*(1,29) = 0.01, *d*_*z*_ = 0.02, *p* = 0.91; Fig 2C]. Follow-up *t*-tests indicated TMR benefitted memory in the high-reward condition [cued: −10.1 ± 7.0 pixels, uncued: 5.3 ± 9.3, *t*(29) = 2.6, *d*_*z*_ = 0.48, *p* = 0.01] and marginally in the low-reward condition [cued: 7.7 ± 5.8, uncued: 22.1 ± 6.9, *t*(29) = 2.0, *d*_*z*_ = 0.37, *p* = 0.052]. Conversely, for CPL, high-reward items were better remembered than low-reward items [*F*(1,29) = 8.9, *d*_*z*_ = 0.55, *p* = 0.006], but there was no effect of TMR [*F*(1,29) = 1.7, *d*_*z*_ = 0.24, *p* = 0.20] or interaction [*F*(1,29) = 0.03, *d*_*z*_ = 0.03, *p* = 0.86; in pixels, high cued: - 3.3 ± 8.0, high uncued: −10.1 ± 9.4, low cued: 14.7 ± 8.9, low uncued: 5.6 ± 7.2; Fig 2D].

### Under separate pair learning, cueing tended to help one item, but not both

Under SPL, when there is less competition during learning, TMR benefitted both high- and low-reward items. However, the above analyses did not examine whether improvements for one item occurred independently of effects on its paired item. For instance, a TMR benefit for Brad Pitt’s spatial location could tend to occur along with a TMR benefit for the Eiffel Tower. If improvements for one item increase the likelihood of improvement for another item, their fates converge; conversely, if it decreases that likelihood, their fates diverge. To test whether their fates converged or diverged, we computed Pearson correlations on the amount of forgetting for all pairs, separately for the cued and uncued conditions within each subject (Fig 3A, top, shows data from one representative subject). Then, we computed paired *t*-tests on Fisher Z-transformed *r* values. We found cued-pair forgetting was significantly less correlated than uncued-pair forgetting [*r*: −0.06 ± 0.04 and 0.11 ± 0.05, respectively: *t*(29) = 2.44, *d*_z_ = 0.45, *p* = 0.02], suggesting more divergence in cued relative to uncued pairs. Further tests revealed uncued-pair correlations were significantly greater than zero [*t*(29) = 2.37, *d*_z_ = 0.43, *p* = 0.02], whereas cued-pair correlations were not—they were numerically (but not significantly) less than zero [*t*(29) = 1.39, *d*_z_ = 0.25, *p* = 0.17; Fig 3A, bottom]. Therefore, cueing seemed to interfere with the positive relationship (i.e., convergence) between items that was present for uncued pairs.

**Figure 3.**
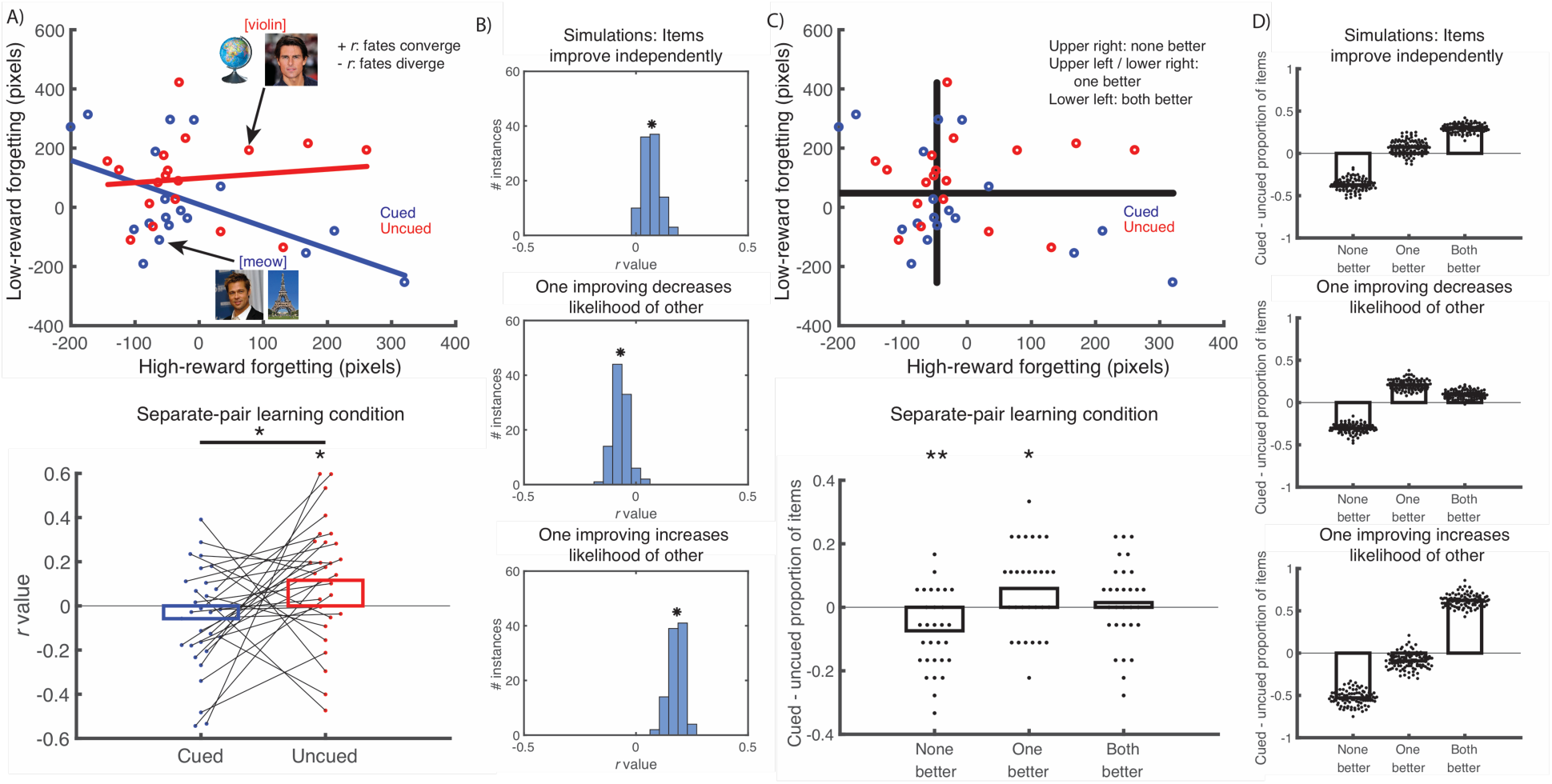
Under separate pair learning, the fate of items within a cued pair diverge, as cueing helps one item, but not both. (A) (upper) We plotted forgetting for each pair of items associated with the same sound and found correlations among cued and uncued pairs. The figure illustrates a single subject’s data. (lower) The fates of cued pairs diverged relative to uncued pairs. Note that statistics were computed on Fisher Z-transformed correlation (*r*) values. (B) We ran simulations of correlations between forgetting of paired items, under different assumptions of how cues affect items within pairs. Under the assumption that cues benefited both items of a pair independently, the positive correlation observed in the uncued condition was carried over into the cued condition (top). Under the assumption that benefits of cueing for one item made the other item less likely to benefit from cueing, correlations became more negative. Under the assumption that benefit of cueing for one item made the other item more likely to benefit, correlations became more positive. Our actual results fit best with the assumption that cueing effects are negatively correlated within a pair. (C) (upper) We computed median forgetting scores for high and low reward items (horizontal and vertical lines) for each subject and calculated the number of pairs in each quadrant (quadrants were defined by each of the paired items being above or below the median forgetting score). If a pair fell in the upper-right quadrant, that indicated that neither of the items in the pair were better than the median *(None better)*, pairs in the upper left and lower right indicated one of the items was better than the median *(One better)*, and pairs in the lower left indicated both items were better (*Both better*). (lower) Plotted are the proportion of cued pairs minus uncued pairs falling within each group defined above. Across subjects, there were significantly fewer cued than uncued pairs in the None better group and significantly more cued than uncued pairs in the One better group. *: *p* < = 0.05. **: *p* < 0.01. (D) Simulation results for the quadrant analysis (under the same three assumptions as above) show, again, that the actual results fit best with the assumption that cueing effects are negatively correlated within a pair.

We next ran simulations (see Methods; Fig 3B) on the data under three assumptions: improvements for one item (1) do not affect the likelihood the other item improves, (2) increase the likelihood the other improves, or (3) decrease the likelihood the other improves. The simulations involved treating the uncued data from the SPL condition as a “baseline” against which cueing could impact results under the various assumptions. The simulations showed that results were most consistent with the third assumption, that a TMR-based improvement in one member of a pair decreased the likelihood the other improved.

In a related analysis, we calculated the median forgetting value for high- and low-reward items separately. Each pair fell into one of four quadrants depending on whether the high-reward and low-reward items were above or below their respective median forgetting values (Fig 3C). Based on this, each pair can be labeled according to whether neither, one, or both items in the pair were higher than the median forgetting value. If cueing improved one item and not the other, we would expect fewer items in the upper right (*None better*) and more in the upper left and lower right quadrants (*One better*), whereas if it improves both items of a pair, there should be fewer pairs in the upper right and more in the lower left *(Both better)*. We found there were significantly fewer cued than uncued pairs in the None better group [*t*(29) = 3.3, *d*_z_ = 0.61, *p* = 0.002], significantly more cued than uncued pairs in the One better group [*t*(29) = 2.39, *d*_z_ = 0.44, *p* = 0.02], and no difference than in the Both better group [*t*(29) = 0.63, *d*_z_ = 0.11, *p* = 0.53]. We ran another simulation to explore what patterns of data we would expect for this quadrant analysis, under different assumptions about how cueing affects paired items. As with our previous simulations, these simulations showed that our results were most consistent with the assumption that cueing improvements are negatively correlated within a pair: If a cue helps one item of a pair, the other item is less likely to benefit from cueing (Fig 3D). Neither the correlation [cued *r*: 0.04 ± 0.04, uncued *r*: 0.05 ± 0.04, *t*(29) = 0.19, *d*_z_ = 0.03, *p* = 0.85] or quadrant analyses produced significant results in the CPL condition [neither cued: 0.27 ± 0.02, uncued: 0.24 ± 0.02, *t*(29) = 1.08, *d*_z_ = 0.20, *p* = 0.29; one cued: 0.48 ± 0.03, uncued: 0.50 ± 0.02, *t*(29) = 0.71, *d*_z_ = 0.13, *p* = 0.49; both cued: 0.25 ± 0.02, uncued: 0.26 ± 0.01, *t*(29) = 0.51, *d*_z_ = 0.09, *p* = 0.62; Fig S2]. Altogether, even for SPL, when we see cueing benefits for both high- and low-reward items, cues do not benefit both memories simultaneously. Instead, each particular cue may be biased to reactivate the high- or low-reward item, thus decreasing the likelihood the other member will also receive benefits.

### Under competitive pair learning, cueing impaired memory for well-learned items

Contrary to what was observed under SPL, we found no cueing benefit under CPL, suggesting competition negatively affects TMR efficacy. Following similar logic, TMR may actually impair memory when competition is strongest, which could occur when both an item and its competitor have strong initial pre-nap accuracy. Additionally, competition might impair low-reward information more than high-reward information, given that subjects had weaker initial memory for low-reward information (Fig 4A).

**Figure 4.**
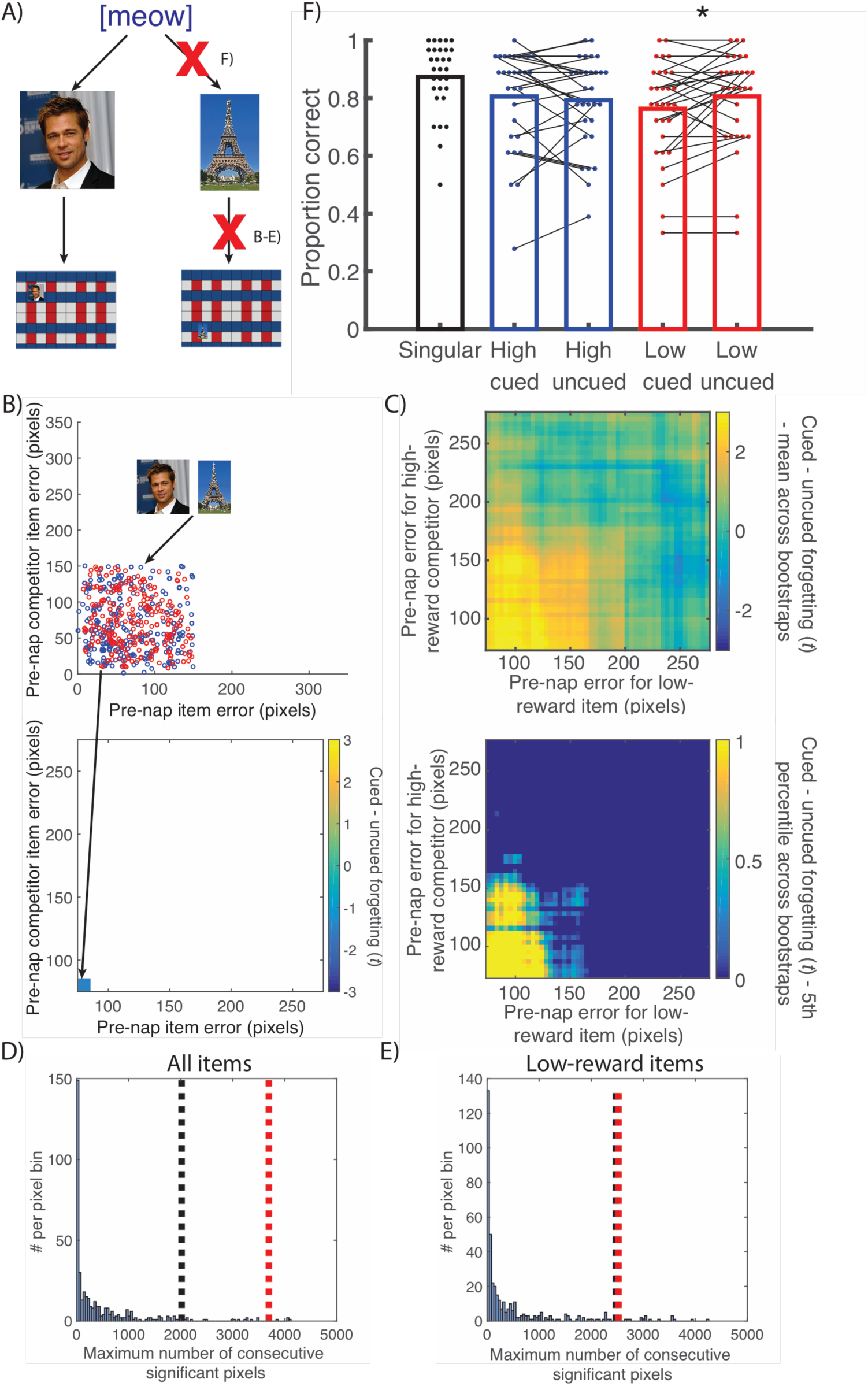
Under competitive pair learning, cueing impairs spatial memory for low-reward items when both items are well-learned pre-nap and also impairs sound-item memory. (A) Schematic showing two ways cueing could impair low reward memories: by weakening associations between the sound and picture later assigned a low reward or the spatial location of the low reward picture item. (B) Schematic of analysis relating pre-nap accuracy to cueing effects. We binned items according to their pre-nap accuracy and the pre-nap accuracy of their competitor. Each bin contained all cued and uncued forgetting values within a moving window (top), then we calculated the *t*-statistic between them (bottom). (C) Using a bootstrap analysis, we found a large cluster of bins that showed a negative TMR effect (more forgetting for cued than uncued items) in the CPL condition; this negative TMR effect is evident for items where both the item itself and its paired item (competitor) were well-learned prior to the nap. The upper plot shows the mean cued-uncued difference in forgetting of low-reward items across bootstraps (results for high-reward items are not shown here). The lower plot highlights only bins where the lower 5^th^ percentile of bootstraps fall above zero. (D-E) Cluster size (red line) exceeds size expected due to randomly shuffling the labels (black line) when combining high reward items against their low reward competitors and low reward items against their high reward competitors (D), as well as low reward items considered against their high reward competitors alone (E). (F) Cueing impaired recall of previously overlearned sound-item associations for items that were assigned to the low-reward condition. * indicates *p* < 0.05.

To test whether TMR impaired memory when competition was strongest, we investigated whether retention differed for cued and uncued items as a function of both the pre-nap accuracy of an item and its competitor. First, for each paired item, we took the item’s pre-nap accuracy and its competitor’s pre-nap accuracy, and we plotted this as a point in 2-d space. We then slid a 150 × 150 pixel moving window around this space (Fig 4B, left). For instance, the bin for 0-150 pixels for an item and 0-150 pixels for its competitor would encompass the pair of Brad Pitt and Eiffel Tower if they had pre-nap errors of 70 pixels and 145 pixels, respectively. Note that, if a pair falls in the lower-left region of this 2-d space, this indicates that both the item and its competitor were well-learned prior to the nap. Next, for each bin (i.e., each location of the moving window), we gathered up all of the items (both cued and uncued) that fell into this region of the 2-d space, and we used a *t*-test to compare the amount of forgetting for cued versus uncued items falling within this bin (Fig 4B, bottom). We repeatedly moved this window until we had covered the entire space. We then repeated these calculations 400 times after resampling subjects with replacement (bootstrapping), producing 400 different *t* values for each bin. We calculated the mean and the 5^th^ and 95^th^ percentiles for the bootstrapped distribution of each bin (Fig 4C); if the bootstrap distribution of *t* values reliably differs from zero, this indicates that the cued-uncued difference is reliable for that bin. Next, we identified clusters of contiguous bins that reliably differed from zero (i.e., the middle 90% of the distribution from the 5^th^ to 95^th^ percentile is above or below zero). Finally, we repeated this entire procedure 400 times after randomly rescrambling cued and uncued labels to find a null distribution of cluster sizes. This allowed us to determine whether the true cluster size exceeded the cluster size expected due to chance (using a *p* < 0.05 family-wise error threshold).

Our first analysis combined both high- and low-reward items. Each pair contributed twice to this analysis: once with the high-reward item as the item of interest (i.e., the item whose forgetting was measured) and once with the low-reward item as the item of interest. In other words, both items acted in turn as the item and the competitor. This analysis produced a significant cluster indicating a TMR impairment in the range in which both an item and its competitor were well-remembered pre-nap (>99^th^ percentile; Fig 4D). We next looked at high-reward items and low-reward items separately. When high-reward items were considered against their low-reward competitor, we found no significant effect (76^th^ percentile), but when low-reward items were considered against their high-reward competitor, we found a significant cluster indicating cueing impairments (>95^th^ percentile; Fig 4E). The same analyses applied to the SPL condition showed a broad range of bins for which cueing was beneficial for memory, meaning it was broadly in the opposite direction of the CPL effect above, but there was no significant cluster (Fig S3). Together, these results demonstrate that under conditions of competitive learning and strong initial memory for more than one item, TMR can cause forgetting.

### Under competitive pair learning, cueing impaired overlearned sound-item memories

After the final post-nap spatial test, subjects took another test to verify that they still retained the sound-item associations learned in Phase 1. We expected recall to be at or near perfect, but also included a *post-hoc* analysis of post-nap scores (not of pre-post differences, because no sound-item test was given prior to the nap). Consistent with the idea that cues linked with more items endure more interference, singular associations were better remembered than all other categories under CPL (proportion correct for singular items: 0.87 ± 0.02, all *p* < 0.005; Fig 4F, on the upper right of Figure 4). However, similar to the cueing impairment in spatial memory for low-reward items under CPL, cues impaired sound-item memory for low-reward items [cued: 0.76 ± 0.03, uncued: 0.81 ± 0.03, *t*(29) = 2.22, *d*_z_ = 0.41, *p* = 0.03] but not high-reward items [cued: 0.81 ± 0.03, uncued: 0.79 ± 0.03, *d*_z_ = 0.12, *p* = 0.52]; the difference in cueing impairment for low-versus high-reward items was marginal [F(1,29) = 4.0, p = 0.055]. Under the SPL condition, we also found better memory for the singular category (proportion correct for singular items: 0.88 ± 0.02, all *p* < 0.005), but found no other effects [high cued: 0.79 ± 0.03, high uncued: 0.79 ± 0.03, *t*(29) = 0.14, *d*_z_ = 0.02, *p* = 0.88; low cued: 0.80 ± 0.02, low uncued: 0.81 ± 0.03, *t*(29) = 0.48, *d*_z_ = 0.08, *p* = 0.63; interaction: *F*(1,29) = 0.05, *p* = 0.83]. Thus, these tests provided converging evidence that low-reward information was weakened under CPL.

### Post-cue beta power negatively predicted subsequent memory and competition-based weakening

Based on previous studies implicating beta power in competition (Waldhauser, Johansson, & Hanslmayr, 2012), we investigated whether post-cue beta power was modulated by competition. Although we expected less competition in the SPL condition than the CPL condition, we expected *both* of the paired conditions to elicit more competition than the singular condition. Thus, we used the difference between singular versus paired to identify competition-sensitive neural signals. In both the CPL and SPL conditions, across-subject differences revealed lower post-cue beta power for singular than paired sounds 250 – 750 ms across multiple electrodes, maximal over electrode FCz in each condition (Fig 5A). We initially used this time segment because it was significant over nearly every electrode. To establish that this difference in beta power was genuine (i.e., not just an artifact of multiple comparisons), we re-confirmed it using a bootstrapping analysis on beta power at FCz. We calculated singular – paired values for each bootstrap. Next, we found clusters of consecutive time points whereby the central 90% of the bootstraps (5^th^ or 95^th^ percentile) differed from zero. Lastly, we computed a null distribution over cluster sizes by repeatedly permuting the conditions across items, re-running the bootstrap, and recording the maximum cluster size for each permutation. We considered a cluster to be significant if its size exceeded 95% of the null distribution (corresponding to a family-wise error rate of .05). Indeed, singular cues had less post-cue beta power than paired cues around the same early beta interval (in relation to the size of clusters from the random (null) distribution for each condition separately: SPL: 97th percentile; CPL: 94.5th percentile; Fig 5A).

**Figure 5.**
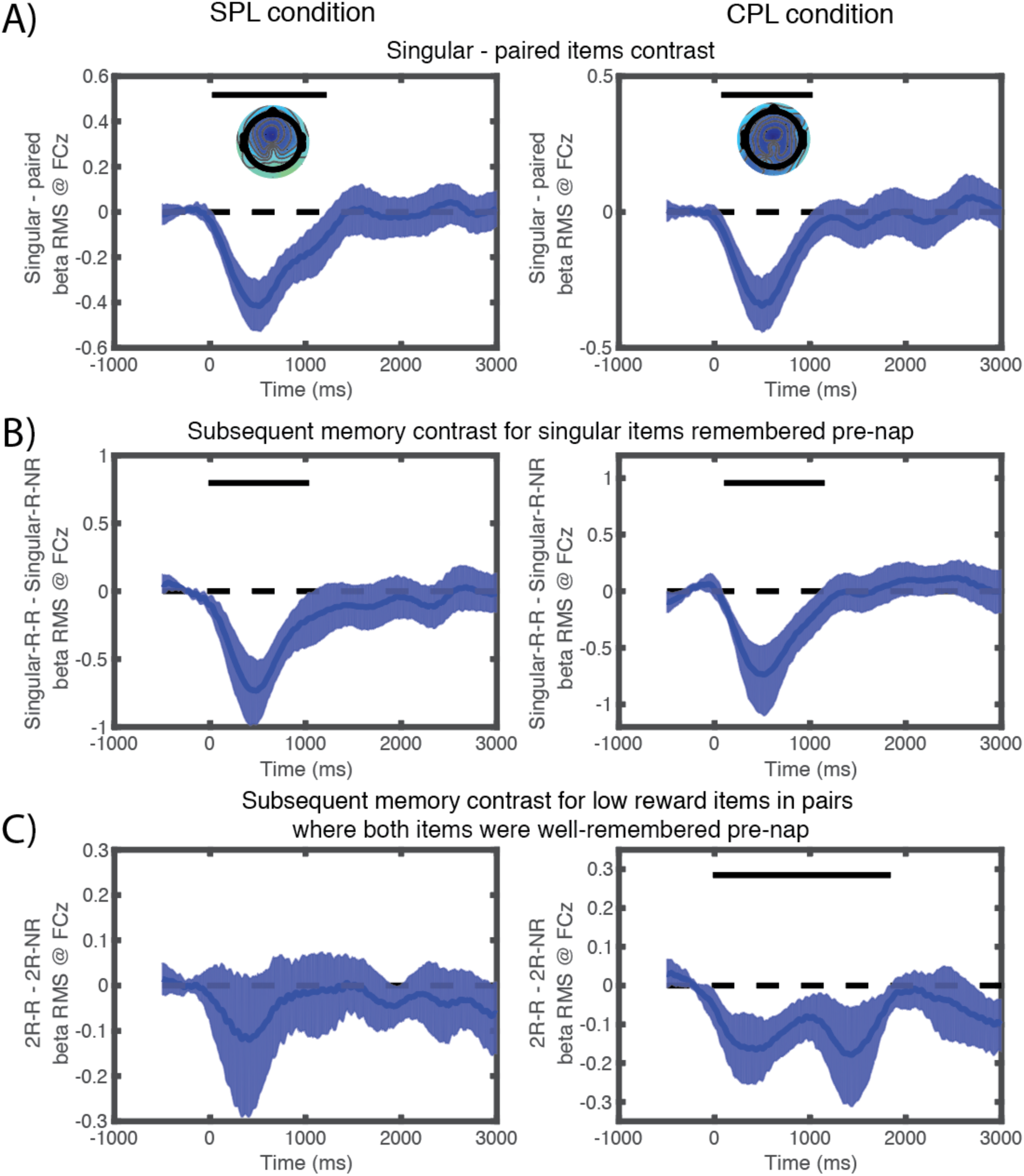
Post-cue beta oscillations increase with competition and negatively predict subsequent memory. (A-C) These plots show the mean surrounded by the central 90% of bootstraps for difference scores of various conditions. (A) Contrast of beta power for singular versus paired sounds. Early post-cue beta power was significantly lower for singular than paired sounds in the SPL (left) and CPL (right) conditions. (B) Contrast of beta power for subsequently remembered versus forgotten items, focusing on singular items that were well-remembered pre-nap. Early beta power negatively predicted subsequent memory for singular items in both the SPL and CPL conditions. (C) Contrast of beta power for subsequently remembered versus forgotten items, focusing on low-reward paired items where both items in the pair were well-remembered pre-nap. Early beta negatively predicted subsequent memory in the CPL, but not the SPL, condition.

Beta power enhancements have also been shown in paradigms where TMR cues impair memory (Oyarzún et al., 2017). One possible interpretation of this finding is that beta indicates competition, which leads to memory weakening. A key prediction that follows from this view is that beta power over FCz (our hypothesized neural indicator of competition) should negatively predict subsequent memory. To test this, we began by investigating the items most likely to become reactivated: those with singular sounds that were well-remembered before the nap (<= 200 pixel error, see Methods), a condition we refer to as Singular-R. We expected competition for these items to be low on average, but we also expected there to be variance across items in the level of competition – singular items can compete to different degrees with other studied items, even if those items were linked to other sounds; we hypothesized that this item-by-item variance would be registered in beta power and would predict memory. Specifically, we asked whether beta power negatively predicted whether these items remained well-remembered or not after the nap (termed Singular-R-R for well-remembered after the nap and Singular-R-NR for not remembered after the nap, respectively) in both CPL and SPL conditions. Individuals without any trials in the Singular-R-NR condition were dropped from the analysis (*N*=2 for SPL, *N*=3 for CPL). Indeed, early beta power was significantly lower for remembered than not remembered items in both conditions (in relation to the size of clusters from the random (null) distribution for each condition separately: SPL: 96^th^ percentile; CPL: 96^th^ percentile; Fig 5B).

Finally, we combined the two ideas about beta power — that it is higher under greater competition and that it predicts memory weakening — to ask how beta influenced the competition-based weakening observed in the CPL condition. Specifically, we asked whether post-cue beta power differed for subsequently remembered versus subsequently forgotten low-reward items from pairs in which both items were initially well-remembered prior to the nap (2R-R [both remembered pre-nap, low-reward remembered post-nap] vs. 2R-NR [both remembered pre-nap, low-reward forgotten post-nap]). Individuals without trials in both conditions were dropped from the analysis (*N*=14 for SPL, *N*=15 for CPL). Indeed, under CPL, post-cue beta power was significantly lower for 2R-R than 2R-NR items (98^th^ percentile). Importantly, this difference was not present in the SPL competition condition, where we did not observe competition-based weakening (53^rd^ percentile). Together, results showed that post-cue beta power increased with competition and negatively predicted memory, including cases of competition-based memory weakening.

### Post-cue sigma power predicted subsequent memory and was reduced under high competition

Based on previous studies, we hypothesized that sigma power approximately 1000-1500 ms post-cue would positively predict retention (Farthouat et al., 2017; Groch et al., 2017; Lehmann et al., 2016; Schreiner et al., 2015). As fast spindles tend to correlate with subsequent memory (James W Antony & Paller, 2017), we chose the midline centroparietal location (CPz) for spindle power *a priori* as it is the scalp location where fast spindle power is maximum (Andrillon et al., 2011; Mölle, Bergmann, Marshall, & Born, 2011; Peter-Derex, Comte, Mauguière, & Salin, 2012). For this analysis, we focused on the same behavioral contrasts as above: first, we looked at singular items that were well-remembered pre-nap and then remembered or forgotten post-nap (Singular-R-R vs Singular-R-NR); next, we looked at low-reward paired items where both items were initially well-remembered pre-nap, and then as a function of whether low-reward items were remembered or forgotten post-nap (2R-R vs. 2R-NR). First, we submitted sigma power to a mixed, condition (SPL vs. CPL) × memory (Singular-R-R vs Singular-R-NR) ANOVA. Individuals without any trials in the Singular-R-NR condition were again dropped from the analysis (*N*=2 for SPL, *N*=3 for CPL). We found a significant effect of memory [*F*(1,53) = 16.8, *p* < 0.001], no main effect of condition [*F*(1,53) = 0.01, *p* = 0.91], and no significant interaction [*F*(1,53) = 0.00, *p* = 0.98]. Follow-up *t*-tests confirmed that sigma power between 1000-1500 ms post-cue was significantly higher for singular items that were subsequently remembered than forgotten in the SPL [mean difference: 0.41 ± 0.14, *t*(27) = 2.86, *d*_*z*_ = 0.54, *p* = 0.008] and CPL conditions [mean difference: 0.41 ± 0.14, *t*(26) = 2.95, *d*_*z*_ = 0.57, *p* = 0.006].

We next asked whether this predictive signal was reduced under conditions in which we found cueing impairments: low-reward paired items when both items were well-remembered. We submitted sigma power to a mixed, condition (SPL vs. CPL) × memory (2R-R vs. 2R-NR) ANOVA. Individuals without trials in both conditions were again dropped from the analysis (*N*=14 for SPL, *N*=15 for CPL). We found a marginally significant main effect of memory [*F*(1,29) = 3.9, *p* = 0.057], no main effect of condition [*F*(1,29) = 0.05, *p* = 0.83], and a significant interaction [*F*(1,29) = 8.6, *p* = 0.007]. Follow-up *t*-tests confirmed that sigma power significantly predicted memory retention in the SPL condition [mean difference: 0.17 ± 0.06, *t*(15) = 2.98, *d*_*z*_ = 0.74, *p* = 0.009], but not in the CPL condition [mean difference: −0.04 ± 0.04, *t*(14) = 0.92, *d*_*z*_ = 0.24, *p* = 0.37]. To look for other time windows that might show an effect, we submitted these analyses to the same bootstrapping procedure that was described above. We found no other time segments showing significance at the 95^th^ percentile. In sum, post-cue sigma power positively predicted memory under low levels of competition, but this signal was less predictive under high levels of competition.

## Discussion

Certain environmental events can cause multiple memories to be activated simultaneously, which produces competition and corresponding changes in memory strength (Lewis-Peacock & Norman, 2014; Norman et al., 2007). Here the amount of competition during learning strongly modulated the effects of TMR during sleep. Under the separate-pair learning (SPL) condition, when the spatial locations of items sharing a common sound were learned separately, TMR improved spatial memory. However, under the competitive-pair learning (CPL) condition, when the locations of items sharing a common sound were learned in succession and rehearsed competitively, TMR produced no overall benefit for memory and even impaired spatial memory when both members of a pair had high pre-nap accuracy (i.e., when competition between the memories was presumably strongest).

Under both learning conditions, high-reward information was retained better than low-reward information. Under SPL, TMR benefited both high- and low-reward information, but further analyses showed that TMR benefits tended to apply to one of the two items within a pair, but not both. Perhaps each cue had a “preferred” item that it preferentially reactivated, and which one was preferred was not necessarily a function of reward value. Future studies that examine inherent preferences for or prior knowledge of each item could provide further clarification.

An entirely different pattern emerged under CPL, whereby TMR impaired sound-item associations that were later assigned a low reward and also impaired spatial memory for low-reward items when both members of a pair were initially well-remembered. Note that the spatial impairment was also observed when considering all items together, but not when considering high-reward items alone. We speculate that weakening occurred because two items simultaneously came to mind, but neither could become fully activated, so each remained only weakly reactivated. This speculation would accord with predictions of the non-monotonic plasticity hypothesis, whereby weak reactivation weakens memory relative to no reactivation at all, whereas only strong reactivation results in strengthening (Lewis-Peacock & Norman, 2014; Newman & Norman, 2010; Norman et al., 2007; Poppenk & Norman, 2014).

Our EEG results align well with a recent model proposing that waking beta power (along with alpha power) plays a crucial role in memory encoding and adjudicating between competing memories at retrieval (Hanslmayr, Staudigl, & Fellner, 2012). As noted earlier, Hanslmayr and colleagues (2009) found that higher beta power at encoding predicted worse subsequent memory. Correspondingly, we found singular items in both learning conditions benefitted from less beta power. Second, Waldhauser and colleagues (2012) found that beta power during wake increased with increased competition between items at retrieval. In keeping with this, we found that TMR cues elicited higher beta power for paired versus singular items in both conditions.

A recent study converged on the idea that beta power may play a role in TMR-based memory impairment (Oyarzún et al., 2017). Participants learned the spatial locations of two identical objects (e.g., dogs; X1-X2 learning) before learning a new location for one of the two objects (X1-X3), followed by sleep and the implementation of TMR. Like our study, this study created a situation in which TMR cues could elicit competition between memories (here, memories of X1-X2 and X1-X3). Critically, X1-X3 learning occurred either immediately after X1-X2 learning (5 min) or after a longer delay (3 hr). Intriguingly, the authors found that TMR improved memory for X1-X2 pairs when learning occurred immediately, but impaired memory when learning occurred with a delay. Furthermore, beta power increased for TMR cues relative to control cues (sounds not linked to studied items), but only in the delayed condition.

These results resemble ours insofar as 1) there was a condition where TMR hurt memory, and 2) these memory weakening effects were associated with beta power increases. However, the results also diverge in important ways. In our study, the condition where there was *no* delay between studying paired items (CPL) yielded memory weakening and beta increases, whereas Oyarzún and colleagues (2017) observed this pattern of results (memory weakening, beta increases) in the delayed condition but *not* the immediate condition. These discrepancies suggest that the lack of a delay in our CPL condition is not, on its own, sufficient to give rise to competition during sleep (see below for additional discussion of this issue). Nonetheless, there were multiple procedural differences between the two studies, and more work is needed to assess which factors are consequential in shaping competitive dynamics during sleep (see Table S2 for a summary of differences between the procedures used in our study vs. those of Oyarzún et al., 2017).

The role of sleep spindles in memory has received substantial support from a vast array of research domains (James W Antony, Gobel, O’Hare, Reber, & Paller, 2012; Bergmann, Mölle, Diedrichs, Born, & Siebner, 2012; Eschenko, Mölle, Born, & Sara, 2006; Latchoumane, Ngo, Born, & Shin, 2017; Mednick et al., 2013; Niknazar, Krishnan, Bazhenov, & Mednick, 2015; Rosanova & Ulrich, 2005). TMR studies have repeatedly shown that sigma (spindle) power approximately 1000-1500 ms post-cue positively predicts memory (Antony, 2015; Farthouat et al., 2017; Groch et al., 2017; Lehmann et al., 2016; Schreiner et al., 2015). Our data from singular items (paired with only a single sound) replicated these findings in both competition conditions. A similar difference was observed in the SPL condition, but was absent from the CPL condition. These results provide further substantiation that post-cue spindle activity benefits memory and that the absence of spindle activity could reflect no or weaker reactivation.

Moreover, results from low-reward items where both of the items in the pair were well-remembered pre-nap demonstrated an intriguing contrast between beta and sigma power. Sigma predicted subsequent memory better in SPL than CPL (Fig 6B), but beta predicted subsequent memory better in CPL than SPL (Fig 5C). Therefore, it appears beta may be more informative than sigma about subsequent memory when competition is high, and vice versa when competition is low.

**Figure 6.**
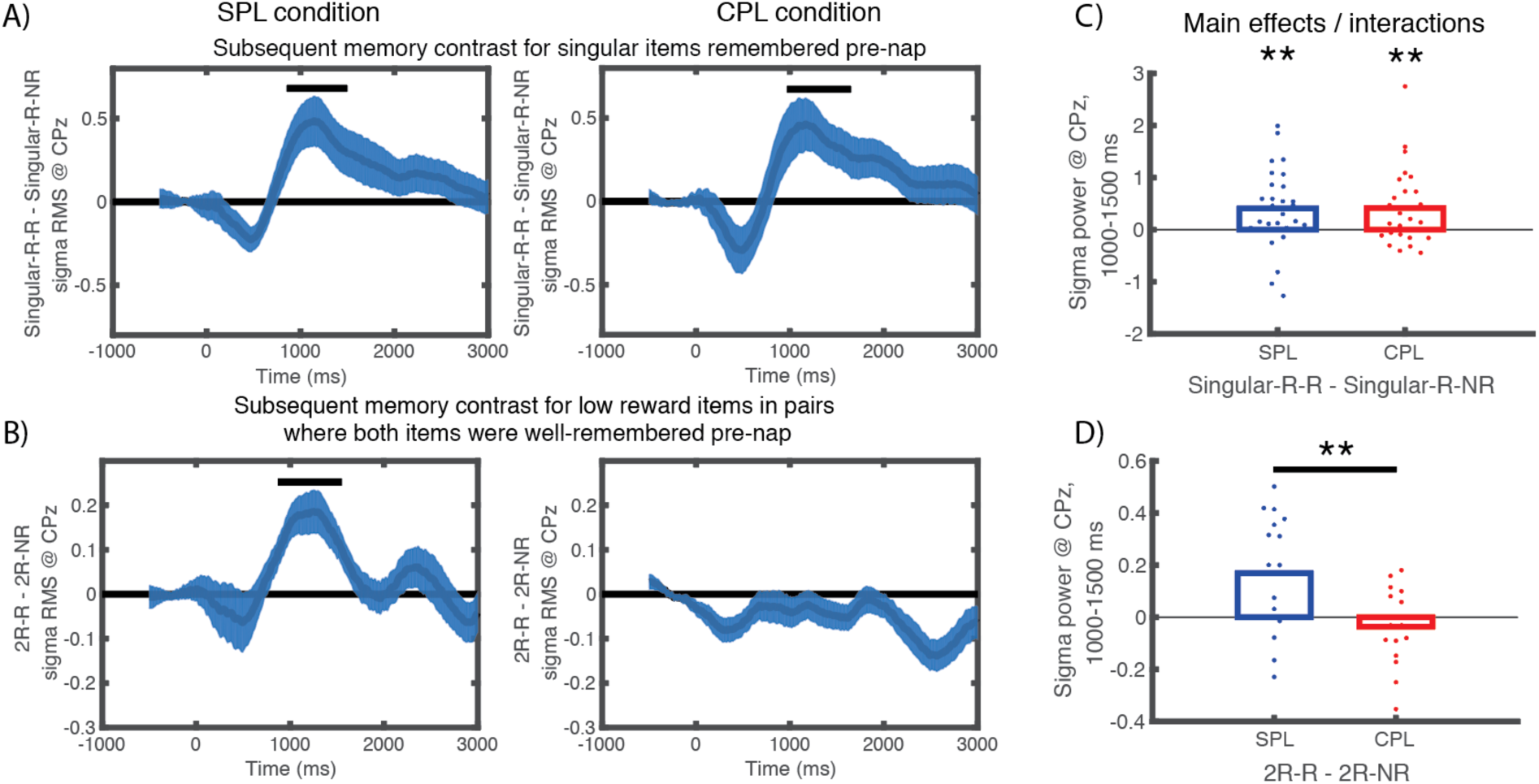
Post-cue sigma positively predict subsequent memory, but not under high competition. We analyzed subsequent memory effects on sigma power for singular items that were well-remembered pre-nap (A) and low reward items within pairs for which both items were well-remembered pre-nap (B). Each plot shows a subtraction of the sigma power trace for items remembered versus forgotten post-nap. (C) For singular items that were well-remembered pre-nap, in both the SPL (left column) and CPL (middle column) conditions, sigma power (1000-1500 ms) positively predicted memory (right column). (D) For low-reward paired items where both items in the pair were well-remembered pre-nap, sigma power positively predicted memory in the SPL condition, but not in the CPL condition. Horizontal bars indicate time points significant at the *p* < 0.05 level.

Importantly, while we observed robust differences in TMR effects following CPL versus SPL, the current study cannot pin down exactly which of the procedural differences between CPL and SPL were responsible for the different results in these conditions. One salient difference between the conditions is temporal proximity (paired items were studied one immediately after the other in CPL, whereas they were separated by a delay in SPL). For reasons noted above (in our discussion of Oyarzún et al., 2017), we think that this temporal proximity factor is not decisive. Another possibility is that our use of a competitive-rehearsal procedure (where participants were instructed to prioritize one item over the other item in the pair) was important for giving rise to the memory weakening and beta increase effects that we observed in the CPL condition. For example, it is possible that results could differ if subjects were asked to integrate, rather than prioritize, the items (Richter, Chanales, & Kuhl, 2016). Another difference between CPL and SPL is that each item in the former condition was studied in succession with the same pairmate, whereas a given item in the SPL condition was studied in succession with a variety of other items (each linked to a different sound). If SPL also used consistent pairings (e.g., meow + Brad Pitt was always rehearsed after violin + globe), this might create competition between these memories during sleep, even if the sounds were different. Future studies could include such alternative conditions to disambiguate these possibilities.

In both learning conditions, paired sound cues could theoretically elicit two memories simultaneously, yet it is only under CPL that we witnessed an absence of TMR benefits (or under certain conditions, reversal). We speculate that under the SPL condition, TMR cues may be biased towards one of the two associations formed at different times during learning, but under CPL, the TMR cues cannot dissociate these two. These findings are especially intriguing in light of studies showing TMR cues benefit memory when linked with an entire learning context, rather than specific trials (Diekelmann et al., 2011; Rasch et al., 2007; Rihm, Diekelmann, Born, & Rasch, 2014). Theoretically, TMR context cues could similarly evoke multiple individual memories, yet they generally benefit memory. These differences raise many questions. Do context cues benefit different memories in a probabilistic fashion on each trial? Do benefits occur because context cues are not especially strongly associated with any particular items, thus decreasing the probability that simultaneous reactivation occurs? Are there subtler forms of competition playing out that get masked by an overall beneficial effect? And more broadly, considering that TMR under the SPL condition tended to benefit only one association, what is the bandwidth of memory reactivation during sleep?

In summary, our study demonstrated systematic differences in memory consolidation during sleep as a function of competition during learning. By pairing sound cues with more than one stimulus in conjunction with competitive learning, our design produced new insights into memory consolidation during sleep. The findings support and extend previous evidence on beta and sigma power, which hold promise for continuing efforts to decipher critical neurophysiological mechanisms in memory processing. The study also expands the scope of the sort of memory processing that can be examined during sleep, beyond individual memories, here emphasizing inter-item competition. Finally, although the physiology of sleep and wake differ substantially from each other, including the near-complete absence of cognitive control in the former, the results are consistent with those during wake showing that simultaneous activation of two pieces of information results in weakening (Lewis-Peacock & Norman, 2014).

## Methods

### Subjects

Sixty subjects (43 female, 18-35 years old) were recruited via online scheduling software at Princeton University (*n* = 37, 27 female) and Northwestern University (*n* = 23, 16 female). Data were collected in approximately similar proportions for the two conditions at the two universities (SPL: 18 Princeton, 12 Northwestern; CPL: 19 Princeton, 11 Northwestern). Forty other subjects (17 Princeton, 23 Northwestern) were excluded for not sleeping long enough for at least one round of sleep cues.Subjects were given hourly monetary compensation for participating and small additional increases based on good performance. This experiment comprised either a separate-pair learning (SPL) condition (*n* = 30, 22 female) or a competitive-pair learning (CPL) condition (*n* = 30, 21 female). Written informed consent was obtained in a manner approved by the Princeton and Northwestern University Institutional Review Boards.

### Stimuli

We included 102 visual stimuli from three categories (celebrities, famous landmarks, common objects), updating the set used by Polyn et al. (2005) for cultural relevance (e.g. Justin Bieber instead of Janet Reno). These items became associated with 66 unique sounds (e.g., “meow”) lasting up to 500 ms adapted from those used by Oudiette et al. (2013). During the nap, sleep cues were embedded in constant white noise (∼ 44 dB), resulting in increases of no greater than 5 dB.

### Design and procedure

The experiment comprised five phases (Fig 1). In phase 1, subjects over-learned arbitrary associations between sounds and items. The goal of this phase was to create strong associations that could consistently support sounds reactivating their corresponding associates during sleep. Thirty sounds were uniquely associated with a single exemplar (10 from each category), while the remaining 36 were associated with two items from different categories. Sound-item mappings were randomly shuffled for each subject. Learning proceeded in four blocks of 20 and one block of 22. Each self-initiated trial began with 1 s of a central fixation cross followed by simultaneous auditory presentation of the sound and visual presentation of the item and sound label. The item was shown centrally with the sound label above. Sound labels were included to eliminate ambiguity of sound identities and to facilitate learning. After 2 s, the sound was repeated and the picture label was included below the picture. After one presentation of each item within a block, we tested subjects by simultaneously presenting the auditory sound, the sound label, and the desired visual category (e.g. “celebrity”). Each sound-item association was tested until it was correctly remembered, after which it dropped out. After all associations from a block were recalled, subjects proceeded to the next block. After the fifth block, all associations were tested until subjects retrieved each correctly again, so in total each association was correctly remembered twice.

In phase 2, subjects learned arbitrary associations between items and locations against a background spatial grid. Each singular item was assigned a high reward and each paired item was assigned a high (9 cents) or low reward (1 cent) to be given at the end of the experiment for correct performance on the pre- and post-nap tests. Subjects were told explicitly these rewards would be given, and reward totals were shown to them and given as shown at the end of the experiment. The goal of this phase was to influence the amount of priority subjects assigned to two items corresponding to the same sound. Its basic structure consisted of subjects viewing one (singular) or two (paired) items for 1 s each, followed by a 5-s rehearsal period where we instructed subjects to maximize their score by freely remembering information in such a way as to give them the highest score. We expected participants would selectively rehearse the high-reward item during this period. Subjects were given breaks at intermittent intervals. Subjects viewed all items four times, each in a new random order. Crucially, for the SPL condition, the two consecutive items before the rehearsal period were never linked with the same sound, whereas in the CPL condition, they were always linked with the same sound. In the SPL condition, paired items were randomly assigned new associates in each round with the requirement that they were not associated with the other item with the same sound. Each item was shown with a height of 150 pixels (5.5 cm) centered around a random location between −300 to 300 pixels (−11.1 to 11.1 cm) from the center of the screen. Each item was shown with its reward value in the center of the picture (Fig 1) and the corresponding sound was played, in order to reinforce associations learned in phase 1. For paired associates, we assigned equal distributions of each possible combination of category-category-reward (e.g., celebrity-high reward + landmark-low reward).

In phase 3, subjects took a pre-nap test by dragging each item from the center of the screen to its location. They indicated their spatial recall choice with a mouse click and were given no feedback.

In phase 4, subjects took an afternoon nap in the lab. Upon online indications of SWS, we administered sleep cues once every 4.5 seconds unless they showed arousals. We cued all of the singular sounds and half of the paired sounds up to seven times. After 60 minutes, if subjects had not received sounds, we administered them during indications of stage-2 sleep. After the nap, subjects left the lab for 2.5 hours.

In phase 5, subjects returned to the lab to take a final spatial memory test followed by a final sound-item test in the same manner as in previous phases. Following these tests, subjects were debriefed and compensated for their participation.

### Dependent variables

We used an adjusted forgetting score as our primary dependent variable. Forgetting, calculated as post-nap error – pre-nap error, significantly correlates with pre-nap error. Items with highly accurate pre-nap recall face ceiling effects (e.g. an error of only 2 pixels cannot be improved across the nap by more than 2 pixels) and those with poor pre-nap accuracy follow a regression to the mean (e.g., an incorrectly recalled location, when very distant from the correct location, is likely to be recalled more accurately after the nap, even by chance). Therefore, we calculated the linear relationship between pre-nap score and forgetting (post-nap – pre-nap score) pooled across subjects in the present data (Fig S1). Then we subtracted each forgetting score from the forgetting expected from this linear relationship (i.e., the residual) to produce the adjusted forgetting store used for all reported analyses. We also calculated accuracy on the final sound-item test; specifically, we calculated the proportion of correctly remembered pictures associated with cued and uncued sounds.

### Permutation tests on pre-nap differences and condition interaction

To ensure differences in pre-nap accuracy could not explain TMR differences between the two learning conditions, we randomly resampled 18-28 subjects without replacement in both the CPL and SPL conditions, selecting 100 instances in which the differences in high-reward pre-nap accuracy between the groups was minimal (*t* < 0.5). We chose a range of sample sizes randomly so the algorithm would not repeatedly choose the same sample of subjects. Then we calculated TMR effects for each of these selections.

### Paired forgetting interactions

To assess whether fates of paired items converged or diverged, we correlated forgetting for the high-reward item with forgetting for the low-reward item separately for cued and uncued conditions for each subject. We then contrasted these *r* values for cued and uncued conditions across subjects with a paired *t*-test. Next, we ran simulations to assess how different assumptions about interactions between paired items will affect these correlations. This involved taking the actual data from uncued pairs in the SPL condition and simulating the effects of cueing under different assumptions about how cueing affects memory. We simulated three different possible effects of cueing, whereby a) paired cues could benefit either item independently (e.g. cue fates unrelated), b) improvement of one item decreased the likelihood that the other improved (e.g. cue fates diverged), and c) improvement of one item increased the likelihood the other improved (e.g. cue fates converged). The high- or low-reward item was randomly chosen to be the first item up as a candidate for improvement and its likelihood of improving was set to 1/9. For the three conditions we simulated, improvements either a) did not change the likelihood of the other item improving (independence), b) reduced the likelihood of the other item improving by a factor of 3 (likelihood=1/27), or c) increased the likelihood of the other item improving by a factor of 3 (likelihood=1/3). After implementing these different effects of cueing, we computed Pearson correlations between the items of each pair on 100 simulations. Note that the uncued SPL condition that we used as a starting point showed a positive correlation (*r* = 0.11; see Results). When we assumed that improvement for an item were completely independent from improvement for its paired item, we continued to observe a positive relationship between forgetting effects for paired items (*r* = 0.07 ± 0.004, *p* < 0.001; Fig 3B, top). When we assumed that improvement for one item decreased the likelihood of improvement for its paired item, this decreased the correlation between items (*r* = −0.07 ± 0.004, *p* < 0.001; Fig 3B, middle). Finally, when we assumed that improvement for one item increased the likelihood of improvement for its the paired item, this increased the correlation between items (*r* = 0.19 ± 0.003, *p* < 0.001; Fig 3B, bottom). Note that significance is less instructive for simulations, as the goal is to capture the qualitative nature of the data under various assumptions.

To further probe the relationship between paired items, we conducted median-split analyses on the amount of forgetting for both high- and low-reward items, creating four quadrants wherein item pairs could fall. Next, we asked how many cued and uncued items fell within each quadrant. We simplified the analyses by considering the upper right quadrant to represent pairs in which neither item was better than the median (*None better*), the lower right and upper left quadrants to represent pairs in which one item was better than the median *(One better)*, and the lower left where both items were better than the median *(Both better)*. Finally, we calculated paired *t*-tests between the proportion of cued and uncued items in each bin. Using the same simulated data that were described above, we assessed how different assumptions about pairwise interactions between items affected the quadrant analysis. As with the previous simulations, we found that our actual results were most consistent with simulation results generated under the assumption that cuing benefits were negatively correlated within a pair (i.e., improvement for one item decreased the likelihood of improvement for its paired item). In the simulation, this condition revealed far fewer cued than uncued pairs in the None better group (mean difference in proportion: −0.30 ± 0.006, *p* < 0.001), far more cued than uncued items in the One better group (0.22 ± 0.007, *p* < 0.001), and only slightly more cued than uncued items in the Both better group (0.083 ± 0.004, *p* < 0.001).

### Item versus competitor pre-nap accuracy bootstrapping procedure

We assessed the influence of TMR on forgetting based on two factors: an item’s pre-nap strength and its competitor’s pre-nap strength. Each bin contained all cued and uncued forgetting values within a moving window of 150 pixels (bin ± 75 pixels, step = 4 pixels; Fig 4B). We then calculated the *t* statistic between the amount of forgetting for cued and uncued items within each bin. We repeated this procedure by randomly resampling subjects with replacement (bootstrapping) 400 times. We determined significance in two steps. First, we sorted all 400 bootstraps and identified clusters of contiguous bins that all differed from zero at the 90% confidence level (between the 5^th^ and 95^th^ percentile). Second, we scrambled the cued and uncued labels 400 times and repeated the bootstrapping procedure, finding the largest cluster size in each scrambled permutation to determine a p < .05 threshold for significant cluster size. Any true cluster size exceeding this threshold was deemed significant.

Cluster maps were calculated separately for all items as well as high-reward and low-reward items only. Note that, for analyses featuring all items, each pair is included twice: The high-reward item stands as the item of interest against the low-reward competitor and the low-reward item stands as the item of interest against the high-reward competitor.

### EEG recording and pre-processing

Continuous EEG was recorded during the nap using Ag/AgCl active electrodes (Biosemi ActiveTwo, Amsterdam) in the same fashion at Northwestern and Princeton. Recordings were made at 512 Hz from 64 scalp EEG electrode locations. In addition, a vertical electrooculogram (EOG) electrode was placed next to the right eye, a horizontal EOG electrode was placed under the left eye, and an electromyogram (EMG) electrode was placed on the chin.

EEG data were processed using a combination of internal functions in EEGLAB (Delorme & Makeig, 2004) and custom-written scripts. Data were re-referenced offline to the average signal of the left and right mastoid channels and were down-sampled to 256 Hz. They were high-pass filtered at 0.1 Hz and low-pass filtered at 60 Hz in successive steps. Problematic channels were interpolated using the spherical method.

### Sleep physiological analyses

Sleep stages were determined by an expert scorer according to standard criteria (Rechtschaffen & Kales, 1968). Table S1 shows the breakdown of stages for each condition as well as the number of cues occurring within each stage. Note that sleep-staging rules require assigning stages based on whichever stage is more prevalent within the 30-s epoch, which can result in sounds occurring in stages that were not the intended targets. Artifacts (large movements, blinks, arousals, and rare, large deflections in single channels) during sleep were marked separately in 5-s chunks following sleep staging.

To calculate oscillatory power, we first filtered separate signals into the sigma (11-16 Hz) and beta (16-30 Hz) bands using a two-way, least-squares finite impulse response filter. Next, we calculated a root-mean-square (RMS) value for every time point using a moving window of 200 ms (using values 100 ms before and after each point) for each channel separately (Mölle et al., 2011; Ngo, Martinetz, Born, & Mölle, 2013). We averaged RMS values within each condition for each subject, ignoring artefactual time segments, and calculated across-subject statistics for our planned contrasts of interest.

We used different approaches for analyzing beta and sigma (spindle) power because we had different predictions given the prior literature. For beta power, we chose electrode FCz based on our finding that the difference in beta power for singular versus paired sounds was greatest for this electrode (effectively, we were using singular versus paired sounds as a “competition localizer”). We used beta power from this electrode as a putative index of competition for all of our analyses comparing beta power for subsequently remembered versus forgotten items. Next, because we were agnostic to time segment, we used a bootstrapping procedure to determine contiguous segments of time in which the central 90% of the data differed from zero and calculated the likelihood that a time segment that large could occur by chance using a p < .05 threshold across the whole interval by scrambling the conditions within each subject. Conversely, for sigma power we had a clear prediction for electrode (CPz) and time segment (1000-1500 ms). We additionally ran bootstrap analyses to ask whether any other sigma power segments (outside of 1000-1500 ms) differed from chance. Finally, some EEG analyses were designed to follow up on the finding that TMR impaired memory in the CPL condition when paired items are well-learned prior to the nap. For these analyses, we set a threshold of pre-nap error <= 200 pixels as indicating “good memory,” which corresponds to the lower-left region in Figure 4C that showed memory weakening effects.

## Acknowledgements

This work was supported by the CV Starr fellowship to JWA and the NSF BCS grant 1533511 to KAP and KAN. We thank Elizabeth McDevitt and Eitan Schechtman for comments on early versions of this manuscript.

